# VirMake: a flexible and user-friendly pipeline for viral taxonomic and functional characterisation from shotgun metagenomic sequencing data

**DOI:** 10.1101/2025.02.07.637044

**Authors:** Einar Birkeland, Paula Istvan, Ekaterina Avershina, Torbjørn Rognes, Trine B Rounge

## Abstract

**Background:** Bacteriophages (phages) are recognized as key regulators of microbial communities, and it is pivotal to understand viral ecology, diversity, and evolution. However, identification and characterization of phages from shotgun metagenome sequencing pose unique bioinformatic challenges owing to the inherent complexity and scale of the data.

**Results:** Here, we introduce VirMake, a comprehensive, flexible, and scalable pipeline developed for viral taxonomic and functional analysis of shotgun metagenome data. It enables high-throughput end-to-end processing, assembly, and annotation of metagenomic data, including viral sequence identification, taxonomic and functional annotation, and using a modular framework, allows users to integrate viral analysis at different stages of metagenome data analysis. VirMake can be installed on Linux operating systems that support Conda and is freely available at https://github.com/Rounge-lab/VirMake. Here, we showcase VirMake capabilities using three metagenome datasets of varying sizes (up to 2.9 Tbp), generated from various environments (human gut and water) and using different sample preparation protocols (VLP enrichment and bulk DNA extraction).

**Conclusions:** VirMake enabled robust processing of all datasets, showing viral genome richness, taxonomy and integration status to vary according to sampling environments and laboratory protocols.

## BACKGROUND

Viruses represent the most common biological entity across ecosystems [1], with virtually all branches of the tree of life being susceptible to their infection. The most diverse group of viruses are those infecting prokaryotes, - the bacteriophages. They are in a constant arms race with their microbial hosts, but they also contribute to host evolution through horizontal gene transfer [2], and to the wider ecosystem through nutrient cycling [3].

Viruses are a highly heterogeneous group and only a small fraction of viral diversity has been captured so far [4]. The study of viruses has been greatly facilitated by shotgun metagenome sequencing, resulting in an exponential increase in the number of publications on metagenome-derived viruses in the last decade, and a concomitant explosion of viral genome sequence data. One outcome of this has been the adoption of a genome-centered taxonomy for viruses [5].

Analysis of shotgun metagenomic data for viral characterisation begins with quality control (QC), which includes filtering of sequences based on quality and trimming sequencing adapters. Following QC, short reads are assembled into contigs. Metagenomic assembly, which differs between short and long read technologies [6], is more complex than single-genome assembly due to sequences from multiple organisms with varying abundances, and it requires specialized assemblers.

Downstream of assembly, metagenomic data analysis workflow depends on the kingdom of interest. The key steps for virome analysis are 1) identification of sequences with a viral origin within contigs, 2) dereplication of viral sequences to generate viral operational taxonomic units (vOTUs), 3) taxonomic and functional annotation of identified vOTUs, and 4) generation of vOTU abundance profiles in samples. Due to high variation in genome lengths and arrangements, host gene acquisition, as well as challenges with separating viral genomes from plasmids [7], and scarcity of reference viral databases, viral sequence identification is crucial for capturing viral diversity. This step is commonly performed using specialized tools employing machine learning algorithms trained on either nucleotide- or protein-based viral signatures [8–10]. Dereplication is performed using algorithms commonly utilized for other kingdoms. This step is computationally expensive and algorithms that perform well with smaller datasets might not be suitable for large metagenomic datasets comprising hundreds of samples. Unlike bacteria, where whole sequence homology search or marker genes identification is employed for taxonomy classification, taxonomy assignment of viruses is often performed based on the gene or protein content [11]. Functional annotation is performed both against general (Pfam [12], KEGG [13]) and specific viral (VOGDB [14]) databases, and additionally includes annotation of the auxiliary metabolic genes (AMGs) that viruses use to modulate host metabolism to their own advantage [15].

Analysis of viral genomics is complex, and there is a need for frameworks that provide non-expert users with easily implementable solutions for end-to-end analysis of viruses derived from metagenome data. Several frameworks for viral genomics and community analysis have been developed (Supplementary Table 1). Some of these pipelines, like ViWrap [16] or phaBOX2 [17], require metagenomic assembly to be performed prior to the analysis, whereas others, like VirPipe [18], allow assembly of both Illumina and Nanopore sequencing data, but do not perform viral sequence identification.

**Table 1.**
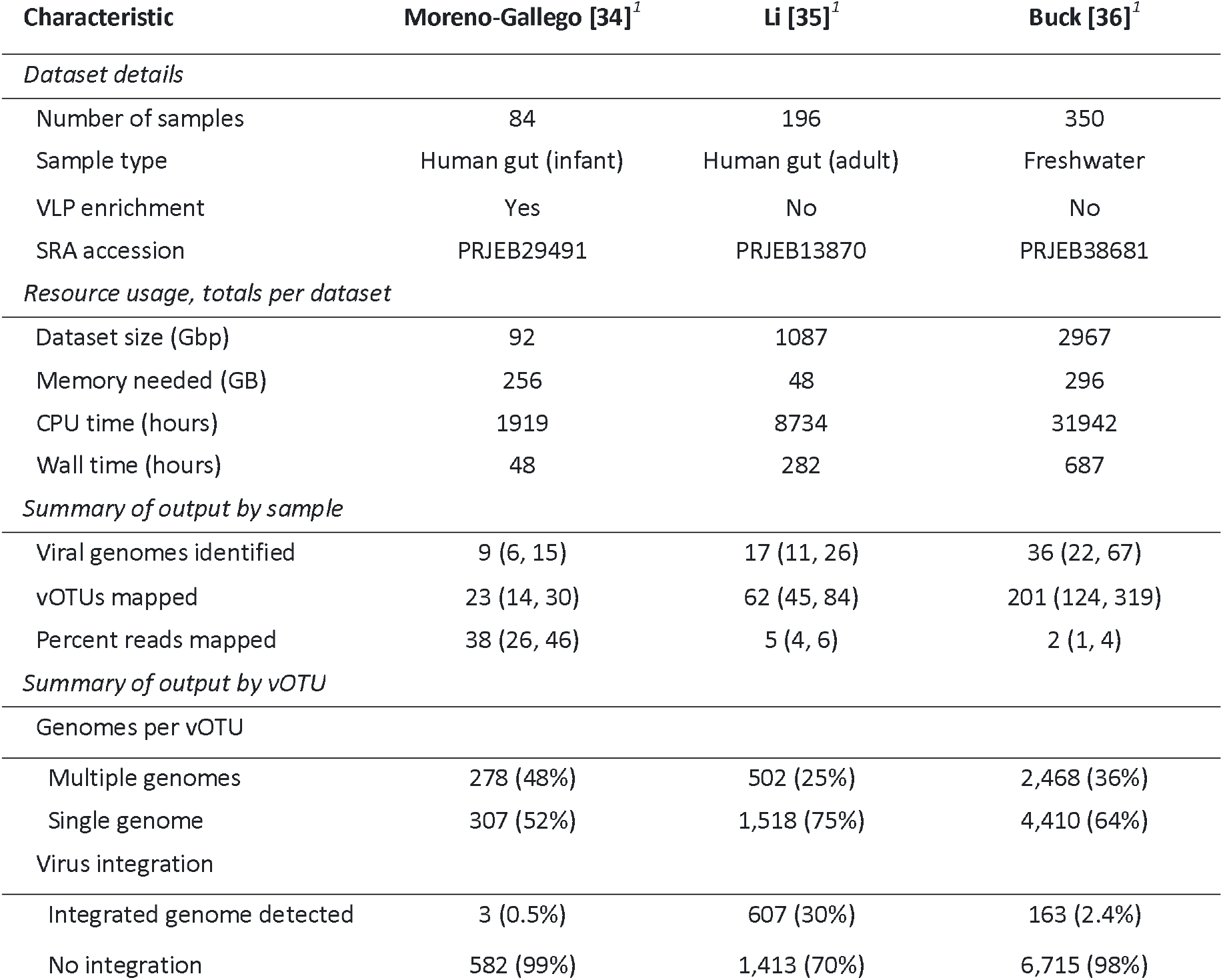
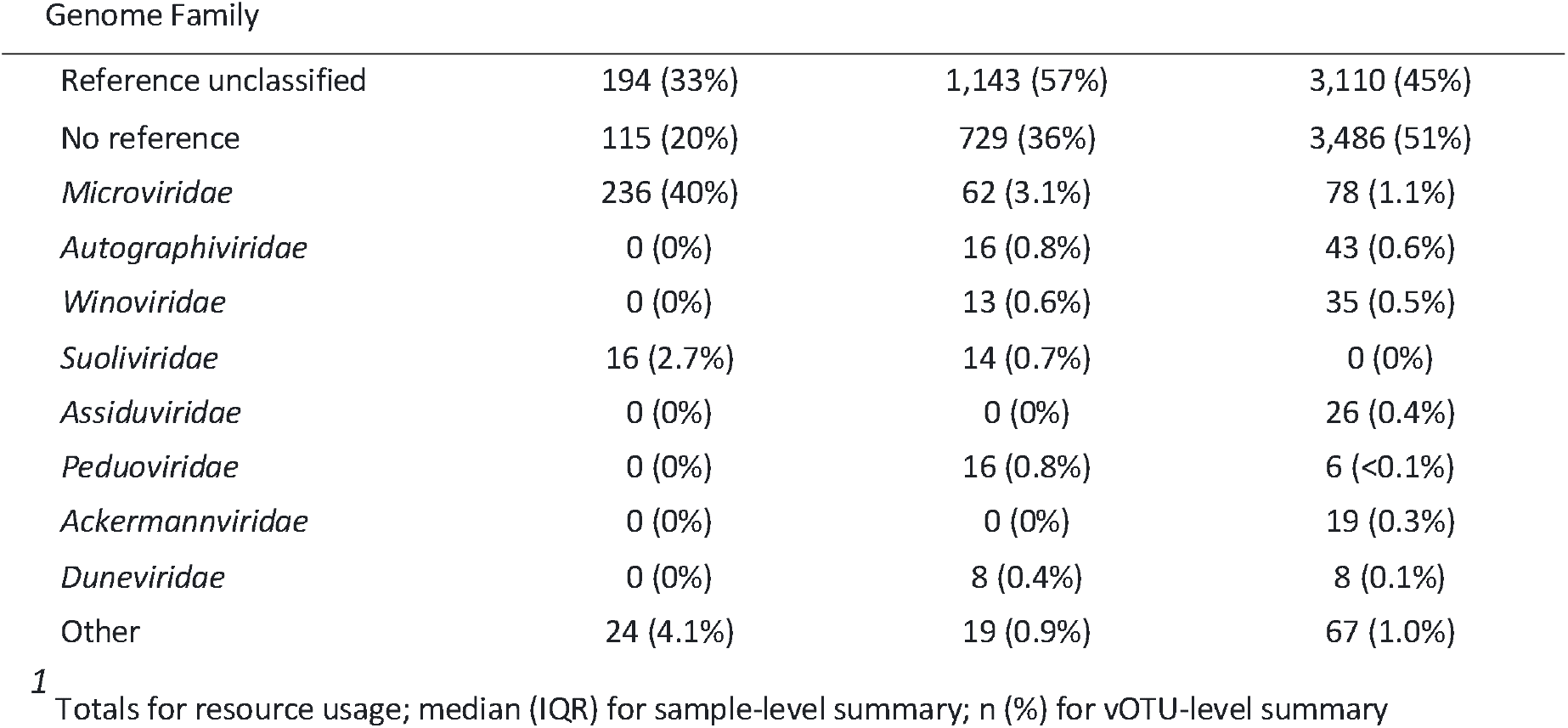
Summary of datasets, computational resources used and results produced by VirMake for the three demonstration datasets.

| Characteristic | Moreno-Gallego [34]<sup>1</sup> | Li [35]<sup>1</sup> | Buck [36]<sup>1</sup> |
| --- | --- | --- | --- |
| *Dataset details* |  |  |  |
| Number of samples | 84 | 196 | 350 |
| Sample type | Human gut (infant) | Human gut (adult) | Freshwater |
| VLP enrichment | Yes | No | No |
| SRA accession | PRJEB29491 | PRJEB13870 | PRJEB38681 |
| *Resource usage, totals per dataset* |  |  |  |
| Dataset size (Gbp) | 92 | 1087 | 2967 |
| Memory needed (GB) | 256 | 48 | 296 |
| CPU time (hours) | 1919 | 8734 | 31942 |
| Wall time (hours) | 48 | 282 | 687 |
| *Summary of output by sample* |  |  |  |
| Viral genomes identified | 9 (6, 15) | 17 (11, 26) | 36 (22, 67) |
| vOTUs mapped | 23 (14, 30) | 62 (45, 84) | 201 (124, 319) |
| Percent reads mapped | 38 (26, 46) | 5 (4, 6) | 2 (1, 4) |
| *Summary of output by vOTU* |  |  |  |
| Genomes per vOTU |  |  |  |
| Multiple genomes | 278 (48%) | 502 (25%) | 2,468 (36%) |
| Single genome | 307 (52%) | 1,518 (75%) | 4,410 (64%) |
| Virus integration |  |  |  |
| Integrated genome detected | 3 (0.5%) | 607 (30%) | 163 (2.4%) |
| No integration | 582 (99%) | 1,413 (70%) | 6,715 (98%) |
| Genome Family |  |  |  |
| Reference unclassified | 194 (33%) | 1,143 (57%) | 3,110 (45%) |
| No reference | 115 (20%) | 729 (36%) | 3,486 (51%) |
| *Microviridae* | 236 (40%) | 62 (3.1%) | 78 (1.1%) |
| *Autographiviridae* | 0 (0%) | 16 (0.8%) | 43 (0.6%) |
| *Winoviridae* | 0 (0%) | 13 (0.6%) | 35 (0.5%) |
| *Suoliviridae* | 16 (2.7%) | 14 (0.7%) | 0 (0%) |
| *Assiduviridae* | 0 (0%) | 0 (0%) | 26 (0.4%) |
| *Peduoviridae* | 0 (0%) | 16 (0.8%) | 6 (<0.1%) |
| *Ackermannviridae* | 0 (0%) | 0 (0%) | 19 (0.3%) |
| *Dunneviridae* | 0 (0%) | 8 (0.4%) | 8 (0.1%) |
| Other | 24 (4.1%) | 19 (0.9%) | 67 (1.0%) |

<sup>1</sup> Totals for resource usage; median (IQR) for sample-level summary; n (%) for vOTU-level summary

Here, we present VirMake, a scalable, efficient, and user-friendly Snakemake-based implementation of viral genomics and community analysis of metagenome sequence data. VirMake is freely available on GitHub: https://github.com/Rounge-lab/VirMake. VirMake is a modular pipeline for sequence data QC and assembly; viral sequence identification and QC; clustering of viral contigs into vOTUs; annotation of viral taxonomy, gene function, and host genome integration; and abundance estimation of vOTUs. Optionally, the user can also assess viral microdiversity. VirMake uses sequencing reads as input, but pre-assembled contigs (from short and long read technologies) can also be provided. Analyses of three short read test datasets showcase VirMake functionalities.

## IMPLEMENTATION

The VirMake pipeline was developed using Snakemake, a workflow management system optimized for scalable and reproducible bioinformatics analyses [19]. VirMake comprises several modular steps including quality control, assembly, clustering, annotation, and classification (Fig. 1). The pipeline is configured for Linux-based environments and supports execution on high-performance computing (HPC) clusters.

**Figure 1.**
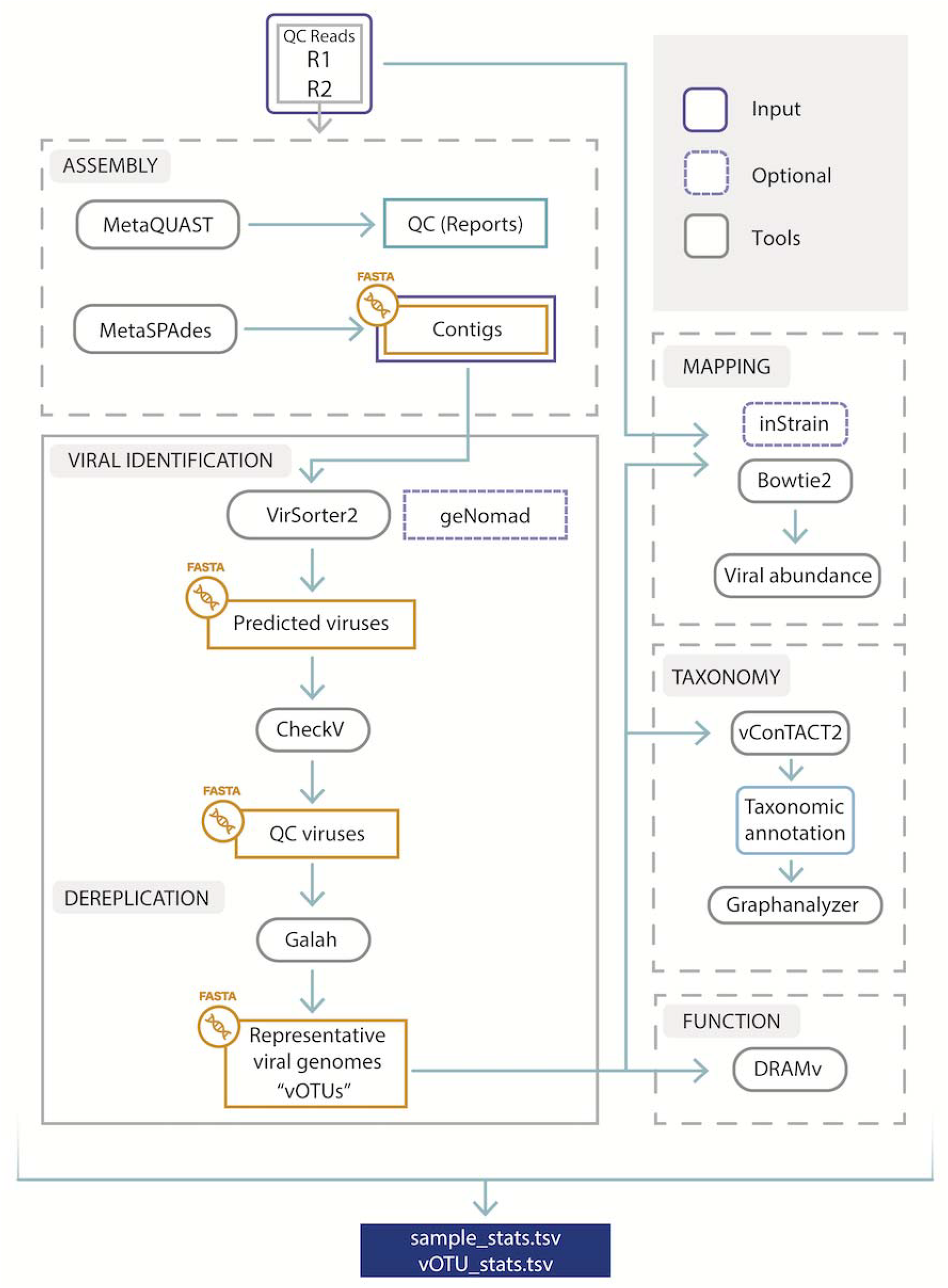
The VirMake pipeline, including modules for i) short read sequencing processing, assembly and quality control, ii) viral identification and dereplication, iii) mapping, iv) taxonomy and v) functional characterization. The pipeline allows users to input (marked in purple) high quality reads in FASTQ format, pre-assembled contigs in FASTA format. Key output variables are presented in two files listing information per sample and representative viral genomes (vOTU).

### Preprocessing, Assembly and QC

VirMake preprocesses paired-end Illumina sequencing data. Initially, the sequencing adapters are removed, and reads are filtered (min base quality = 15, min filtered read length = 15) using fastp [20]. High quality reads are then assembled into contigs using MetaSPAdes [21], an assembler optimized for metagenomic data. The quality of the assembled contigs is evaluated using MetaQUAST [22]. The contigs are then fed into the Viral identification and dereplication module. The user can optionally provide VirMake with pre-assembled contigs as input. In that case, the preprocessing and assembly modules will not be executed.

### Viral identification and dereplication

To identify viral sequences within a preprocessed or preassembled set of contigs, VirMake employs VirSorter2 [10] (default) or geNomad [23]. The user can specify minimum input lengths, with the default set to 1000. CheckV [24] is used for evaluation of viral genome quality and completeness for all contigs with predicted viral sequences, with a user-defined threshold for inclusion in subsequent analyses (default set to minimum of “Medium quality”, i.e. ≥ 50 % completeness). The output is restricted to genomic regions predicted as viral by both the virus identification tool and CheckV. Extracted viral genomes are then clustered into vOTUs using Galah [25] based on a user-specified average nucleotide identity (ANI) threshold (default: 97) across a user-specified fraction of each genome (default: 70). For each vOTU, the viral genome with the highest completeness as determined by CheckV is defined as the vOTU representative.

### Mapping, viral abundance and microdiversity

Bowtie2 [26] is used to map the quality-controlled reads to the representative viral genomes. For abundance estimation, the median depth of coverage is calculated using BBTools’ pileup function [27], with only those vOTUs achieving at least a user-defined (default≥75%) genome coverage considered to have a genome present. An optional strain-level diversity analysis can be conducted by selecting to use inStrain [28], set to determine population-level sequence diversity for each vOTU. Use of inStrain is not enabled by default due to the potentially high resource usage.

### Taxonomy

Taxonomic classification in VirMake is performed using vConTACT2 [11], employing Prodigal [29] for gene identification, and INPHARED [30] for reference genome database construction. Definition of genome co-clustering is conducted using an algorithm developed as part of MetaPhage [31], graphanalyzer (default settings) [32], based on defining closely clustering and neighbouring genomes in the gene sharing network created by vConTACT2.

### Function

Gene annotation of vOTU genomes is incorporated as an optional module in VirMake. Here, DRAM-v [33] is used with the default databases (Pfam, VOGDB, KOfam, dbCAN, and RefSeq), and includes identification of auxiliary metabolic genes (AMGs). Database construction is a resource-demanding procedure that is only carried out if gene annotation is enabled.

### Reports generation

VirMake generates summaries of processed data at the level of samples and vOTUs. These provide an overview of the analyzed viral genomes and their associated metadata. The sample-level statistics file compiles key statistics for each sample, including the total number of viral genomes recovered from contigs, and details on their length distribution, genome quality, and the number of integrated viral sequences. Further information includes the number of vOTUs detected using read mapping, and statistics on assembly quality. The file with information on vOTUs summarizes information from taxonomic classification, and characteristics of the representative genome are provided, including length, genome quality, completeness, contamination, integration state, and a summary of gene annotation. Further information about the pangenome of each vOTU is reported, including the number of viral genomes clustering together with the representative, along with information on length distribution, and rate of genome integration.

### Test datasets

We employed three publicly available metagenome datasets of varying sample numbers, viral enrichment protocols, and sequencing depth (Table 1). Two of these studies targeted the human gut microbiome, and one reported shotgun metagenome sequence data from freshwater reservoirs [34–36]. All datasets were run using default parameter settings, except for the microdiversity profiling module being enabled.

## Results

An overview of the resource usage of VirMake for the three datasets is shown in Table 1 and Figure 2. Metagenome assembly was the most resource demanding step, both in terms of CPU time and memory load (Figure 2). The maximum amount of memory required was 296 GB for one of the jobs of the Buck dataset. Further details are provided in Supplementary Tables 2 and 3. Virus identification was the second most demanding module, although InStrain microdiversity analysis also used considerable amounts of memory, at least with the Li et al. dataset. We attempted benchmarking VirMake against MetaPhage and ViromeFlowX as these pipelines are most similar to VirMake (see supplementary table 1 for overview of pipelines), but their implementation failed due to technical shortcomings (Supplementary Text 1).

**Figure 2.**
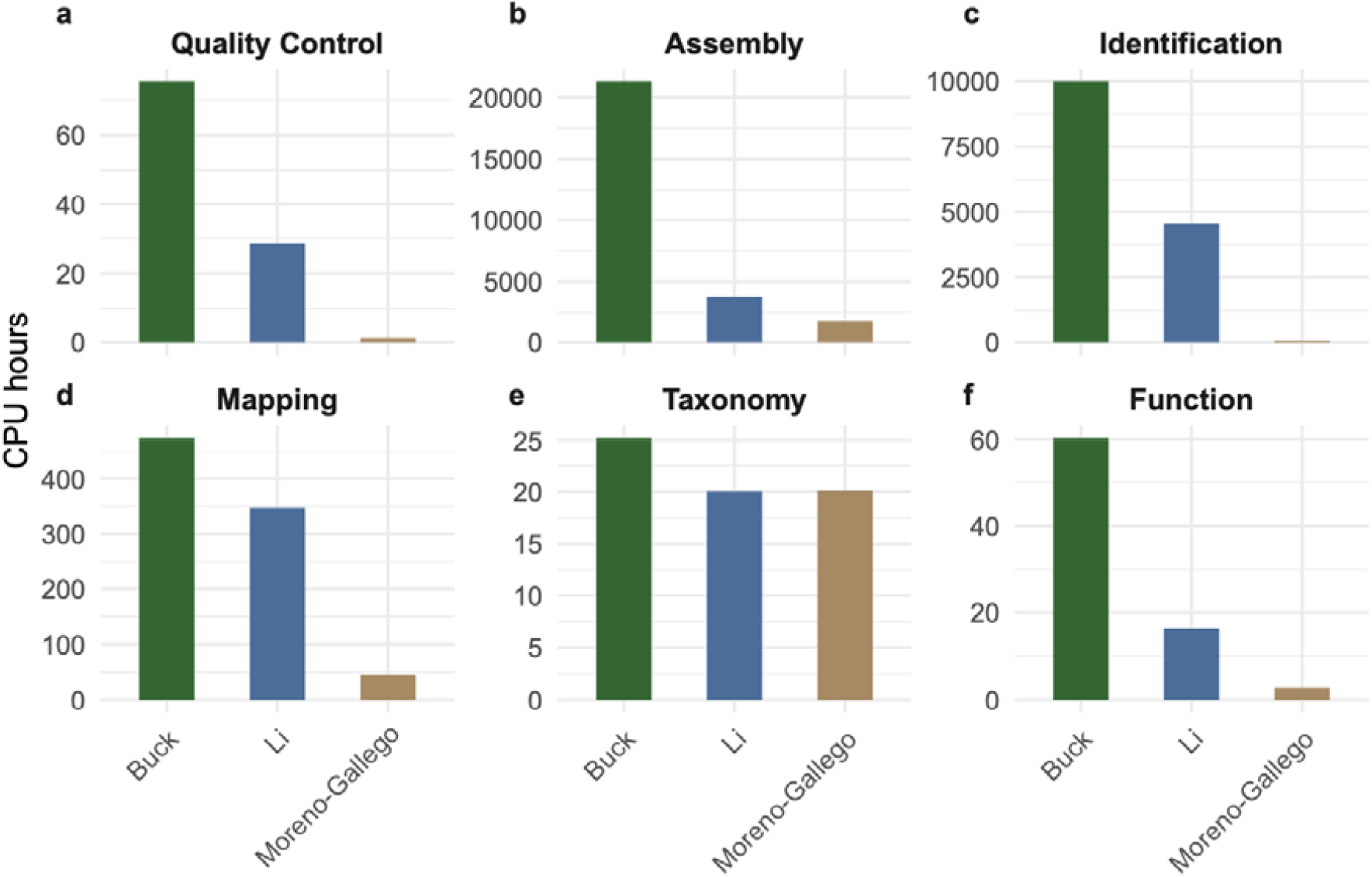
CPU time when running VirMake on three datasets. Included datasets are those published in Buck et al. [36], Li et al. [35], and Moreno-Gallego et al. [34]. The CPU time is provided for each module: a) quality control, b) assembly, c) identification, d) mapping, e) taxonomy, and f) functional annotation.

Using default settings for viral identification, a median of 9, 17 and 36 viral genomes were recovered per sample from assembled contigs in the Moreno-Gallego, Li and Buck datasets, respectively (Table 1). The number of dataset representative viral genomes (vOTUs) detected using read mapping showed the same pattern (median of 23, 62, and 201 vOTUs per sample, respectively). Both the number of recovered genomes and detected vOTUs correlated with sequencing depth and assembly size (Supplementary Figure 1). The median (IQR; dataset) number of genes per vOTU ranged between 11 (42; Moreno-Gallego) and 45 (32; Buck), whereas gene density remained stable across datasets and comprised 1.4 (0.3) genes per 1000 bp of viral sequence. Auxiliary metabolic genes were identified in 84 %, 79 % and 74 % of vOTUs in Moreno-Gallego, Li and Buck datasets, respectively.

Contrasting the samples in the Moreno-Gallego dataset (human gut, infants) to that of bulk metagenome samples in the Li dataset (human gut, adults), we found a clear enrichment of read mapping to vOTUs in the former, likely due to their use of VLP enrichment. Here, most vOTUs were clustered with genomes in the reference database, and as much as 40% were assigned to the *Microviridae* family (Table 1; Figure 3). The overall viral diversity was higher in bulk metagenome sequenced samples (Li), including a higher number of viral genome recovery, a higher rate of single-genome vOTUs, but fewer reads mapping to vOTUs. One reason for this is likely the identification of more viral genomes in a prophage state, as was indicated by the higher rate of genome integration (30% of vOTUs with at least one integrated genome member in Li vs <1% in Moreno-Gallego; Figure 3).

**Figure 3.**
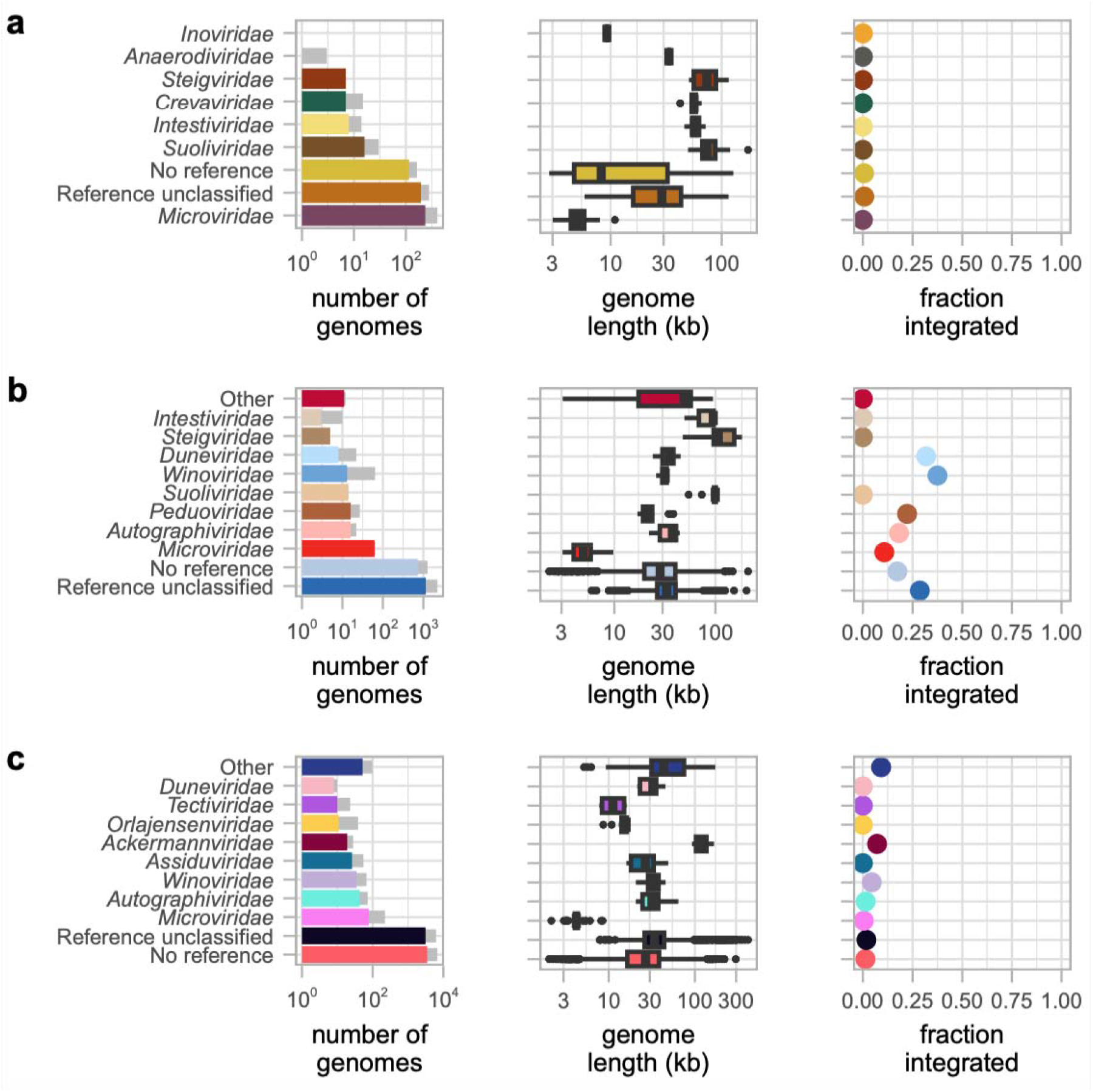
Viral genome characteristics in three metagenome datasets. a-c) left panel: The number of viral vOTUs assigned to each taxonomic family is shown in colored bars, with the total number of viral genomes represented by the vOTUs shown in grey bars; middle panel: Distribution of genome lengths for vOTUs belonging to each taxonomic family; right panel: The fraction of viral genomes determined to be in an integrated state across vOTUs for each viral family is reported. Reference unclassified: query genome clusters with a known reference genome that lacks taxonomy assignment on a family level. No reference: no clustering between the query and any reference genomes. a) characteristics of viral genomes detected in the 84 VLP-enriched gut metagenome samples from Moreno-Gallego et al.[34], including 919 viral genomes, and 585 vOTUs. b) characteristics of viral genomes detected in the 196 gut metagenome bulk sequenced samples from Li et al., including 3938 viral genomes, and 2020 vOTUs. c) characteristics of viral genomes detected in the 350 freshwater lake samples (61 isolated bacteria and 289 metagenomes) from Buck et al., including 13 629 viral genomes, and 6878 vOTUs.

The dataset derived from freshwater samples had fewer reads mapping to vOTUs compared to the human gut bulk metagenome dataset (median 2% vs 5% for Buck and Li datasets respectively), and vOTUs in the Buck dataset were more commonly found to represent multiple viral genomes, but it is possible that this increase was caused by saturation effects, as the Buck dataset was substantially larger than the other two. The Buck dataset also included 61 single amplified genomes (SAGs) of individual bacterial cells. In these samples, the number of identified genomes was lower than in the freshwater microbiome samples (median (IQR) n = 0 (1) vs n = 36 (45) genomes per sample).

## Discussion

We have developed a scalable end-to-end pipeline for the extraction, characterisation and functional analysis of viral sequences from metagenomic datasets. The pipeline is arranged in modules that can be invoked both individually and collectively, using FASTQ files or assembled contigs. Resulting files from each step are provided in corresponding folders and their outputs are summarized in two user-friendly reports for samples and vOTU characteristics.

Viral analysis gains increasing attention and several pipelines for virome analysis exist, however, many of them lack versatility, are not actively maintained, or do not scale to enable analyses of hundreds of samples. Here we show VirMakes capabilities by analysing three different metagenome datasets, with the largest being a 350 sample dataset, comprising 2.9 Tbp of sequencing data. We have also previously employed a beta-version of VirMake in our recent virome analysis of >1000 stool samples [37].

The major challenge with large datasets is dereplication of viral genomes. CD-HIT, implemented in ViromeFlowX [38] or MetaPhage [31], is an established alignment-based algorithm widely used for redundancy reduction [39]. However, its runtime increases quadratically with input size [40]. In VirMake, we have incorporated Galah, - an alignment free tool that uses a greedy clustering algorithm based on average nucleotide identity (ANI) distances to reduce computational time. Alternatively, some pipelines like ViWrap [16] or phageannotator (in development) [41], employ vRhyme [42] for binning of viral genomes instead of dereplication.

Metagenome assembly is another highly resource-demanding step. FastQC quality control (QC) and assembly were included in VirMake to allow for a complete pipeline from sequencing reads to viral characterisation. These steps are optional, allowing for cases where sequencing data have been quality controlled or pre-assembled. In many pipelines, however, assembly is either hardwired, like in ViromeFlowX and MetaPhage, or absent, like in ViWrap [43] or phaBOX2 [17]. To our knowledge, the optional inclusion of data assembly is available only in VirPipe [18] and VirMake.

Comparing VirMake to other pipelines with regards to taxonomy classification, VirPipe and ViromeFlowX employ DNA based homology to reference sequences using Kraken2 [44], whereas VirMake, MetaPhage and ViWrap implement network-based taxonomy inference. Since viruses are polyphyletic and prone to horizontal gene transfer [45], gene-sharing networks based on protein sequences provide better taxonomy resolution [11]. Moreover, viral diversity is poorly represented in reference databases [4] leading to even lower identification rates when using DNA homology. By default, VirMake identifies viral sequences using VirSorter2, which minimizes the error of non-viral circular sequences misclassification to viruses [10], however, geNomad with simultaneous viral and plasmids sequences identification, is also implemented for users preferring this tool.

We have attempted to benchmark VirMake to MetaPhage and ViromeFlowX since these pipelines are very similar to VirMake with regards to processing and output. We failed to benchmark MetaPhage due to Conda packages incompatibility and unresponsive URLs. ViromeFlowX also exhibited several challenges with installation, database retrieval and with running of metaSPAdes and VirFinder, rendering the pipeline challenging to benchmark.

The rapid evolution of viral classification necessitates continuous adaptation to reflect new discoveries and nomenclature changes. Future work should explore the integration of alternative taxonomic classifiers, such as ViraLM [46], to enhance accuracy and broaden applicability.

## Conclusion

VirMake provides a comprehensive and flexible solution for viral taxonomic and functional analysis from shotgun metagenomic data. It supports analyses of large and diverse datasets. Updates will be essential to keep pace with evolving viral taxonomy and analytical advancements, ensuring VirMake remains a valuable tool for the research community.

## Declarations

### Availability of data and material

The source code for VirMake is available at https://github.com/Rounge-lab/VirMake.

The public datasets are available for download at the sequence read archive with accession numbers

PRJEB29491 (Moreno-Gallego), PRJEB13870 (Li), and PRJEB38681 (Buck).

Code to reproduce the analyses presented in the manuscript is available at https://github.com/Rounge-lab/VirMake-demo.

### Authors’ contributions

TBR and TR conceptualized and designed the study and supervised the project. EB, TR and PI developed the software, including its implementation. EB and EA conducted analyses of datasets. All authors drafted the manuscript and approved the final manuscript.

## Acknowledgements

We would like to thank Simon Norvold Barak, who contributed to the initial implementation of VirMake. Computations were performed on resources provided by Sigma2 - the National Infrastructure for High-Performance Computing and Data Storage in Norway and on the Educloud Fox cluster provided by the University of Oslo.

