## Supplementary information for "VirMake: a flexible and user-friendly pipeline for viral taxonomic and functional characterisation from shotgun metagenomic sequencing data"

##### Supplementary Information 1. Running MetaPhage and ViromeFlowX

The MetaPhage pipeline [1] for viral metagenome data analysis was identified as the most similar to VirMake (Supplementary Table 1). We tried to install and run both the main (0.3.3) and development (beta) version on the GitHub repository (https://github.com/MattiaPandolfoVR/MetaPhage; last updated December 2023) of the pipeline for comparison. Due to multiple unresolved issues (including an unresponsive database URL and incompatible Conda packages) this was unsuccessful.

The Nextflow based ViromeFlowX workflow [2] was also identified as a relevant pipeline (Supplementary Table 1). We tried to install the most recent version from the GitHub repository (https://github.com/01life/ViromeFlowX; last updated April 2024) for comparison, but were unsuccessful in running it. After overcoming a few lacking dependencies (the seqtk and parallel Conda packages), the samples could not be processed due to problems arising when running VirFinder [3] and metaSPAdes [4]. A major obstacle was the lack of a script for downloading the required databases in the appropriate folders; for most databases only a set of URLs were provided.

*References:*


1. *Pandolfo M, Telatin A, Lazzari G, Adriaenssens EM, Vitulo N. MetaPhage: an Automated Pipeline for Analyzing, Annotating, and Classifying Bacteriophages in Metagenomics Sequencing Data. mSystems. 2022;7:e00741-22.*
2. *Wang X, Ding Z, Yang Y, Liang L, Sun Y, Hou C, et al. ViromeFlowX: a Comprehensive Nextflow-based Automated Workflow for Mining Viral Genomes from Metagenomic Sequencing Data. Microb Genomics. 2024;10:001202.*
3. *Ren, J., Ahlgren, N.A., Lu, Y.Y. et al. VirFinder: a novel k-mer based tool for identifying viral sequences from assembled metagenomic data. Microbiome 5, 69 (2017).*
4. *Nurk S, Meleshko D, Korobeynikov A, Pevzner PA. metaSPAdes: a new versatile metagenomic assembler. Genome Res. 2017 May;27(5):824-834.*

#### Supplementary figures

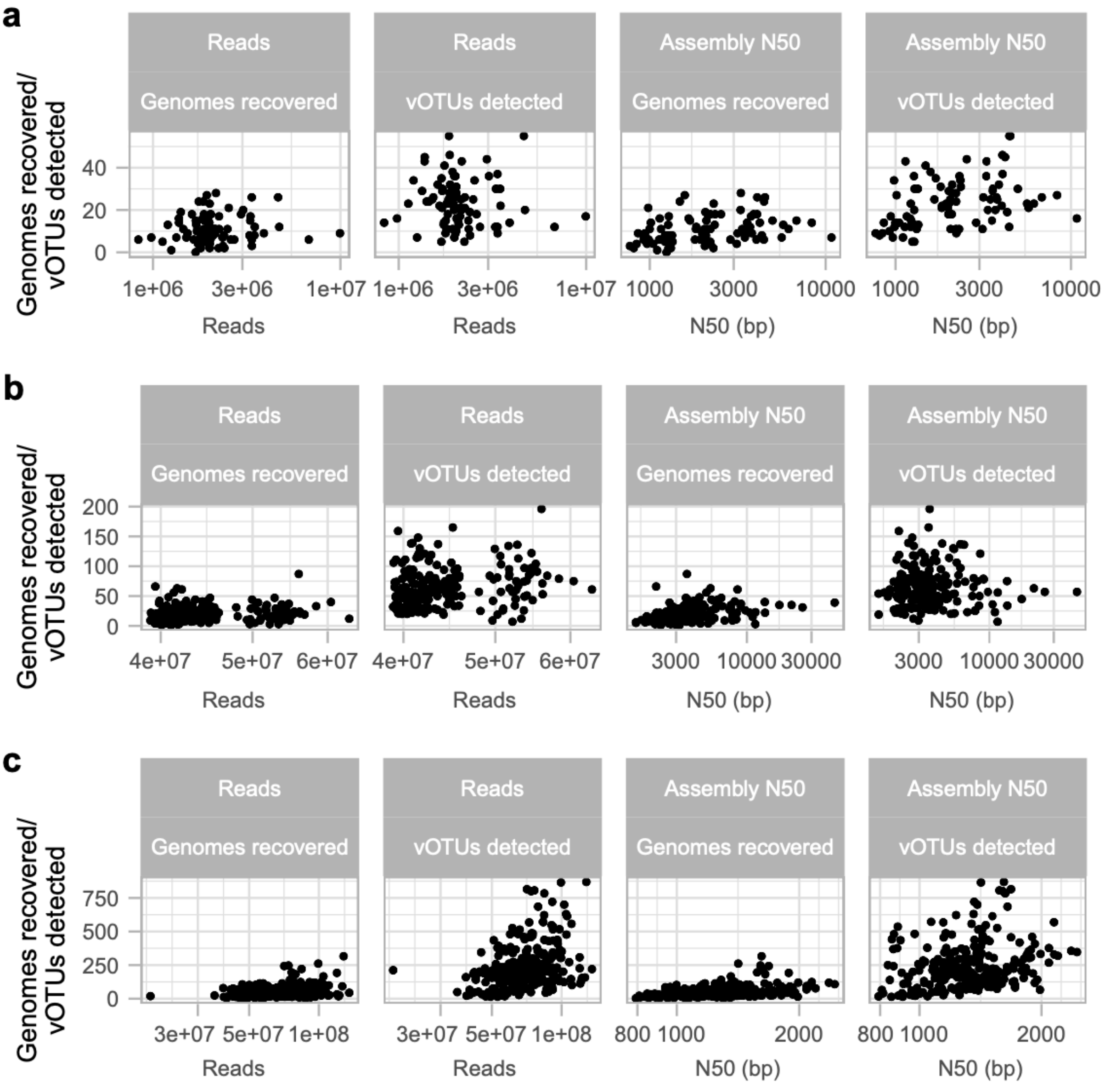


**Supplementary Figure 1. Sequencing data and assembly used as input and number of viruses identified and detected.** a-c) Number of QC sequencing reads (“Reads”) used as input to VirMake, and assembly N50 of contigs prior to viral identification are shown on the x-axis, and the number of genomes recovered using viral identification algorithms (here, VirSorter2), and number of vOTUs detected using read mapping to representative viral sequences (“vOTUs detected”). Results are presented for a) Moreno-Gallego, b) Li, and c) the Buck dataset. Only results derived from metagenome sequencing samples are presented for the Buck dataset.

### Supplementary tables

**Supplementary Table 1. Pipelines for viral data analysis from shotgun metagenomic data**

| Tool | Year | Input type | Assembly | Viral contigs identification | Viral contigs quality check | Dereplication (vOTU clustering) | vOTU mapping | Taxonomy classification | Functional annotation | Link |
|---|---|---|---|---|---|---|---|---|---|---|
| MetaPhage | 2022 | reads (Illumina) | MegaHIT, metaQUAST | DeepVirFinder, Phigaro, VIBRANT, VirFinder, VirSorter2 | - | CD-HIT | bowtie2, BamToCov | Kraken2, vConTACT2 | - | https://github.com/MattiaPandolfoVR/MetaPhage |
| ViWrap | 2023 | contigs | - | VIBRANT, VirSorter2, DeepVirFinder | CheckV | vRhyme, dREP | vRhyme | vContact2, NCBI Refseq, VOG HMM | EggNOG, PROKKA, KEGG/KOfam, VOGDB, Pfam | https://github.com/AnantharamanLab/ViWrap |
| VirPipe | 2023 | reads (Illumina and Nanopore) | SPAdes, MegaHIT, canu, flye | - | Qualimap | - | Minimap2 | Kraken2, Centrifuge | BLASTx/BLASTp, InterProScan, Pfam, COG | https://github.com/KijinKims/VirPipe |
| VIRify | 2023 | contigs | - | VIBRANT, VirSorter2, PPR-Meta | CheckV | - | - | ViPhOGs | VOGDB, Pfam, TIGRFAM, KEGG | https://github.com/EBI-Metagenomics/emg-viral-pipeline |
| ViromeFlowX (nf-core) | 2024 | reads (Illumina) | metaSPAdes | VirSorter2, VirFinder | CheckV | CD-HIT | bowtie2 | Kraken2 | PROKKA, EggNOG, KEGG/KOfam, VOGDB, InterProScan, Pfam | https://github.com/01life/ViromeFlowX |

| | | | | | | | | | | |
|---|---|---|---|---|---|---|---|---|---|---|
| PhaBOX v2.0 | 2024 | contigs | - | PhaMer | PhaMer | PhaBox2 | - | PhaGCN (ICTV 2024, vOTU clustering, protein annotation) | PhaVIP | https://github.com/KennthShang/PhaBOX |
| ViruSpy | 2017 | reads (Illumina) | MegaHIT | BLAST+Glimmer | - | - | - | RefSeq Viral Genome Database | - | https://github.com/NCBI-Hackathons/ViruSpy |
| VIP2 | 2019 | reads (Illumina) | Velvet-Oases | BLAST | - | - | bowtie2 | Kraken2, Kaiju | - | https://github.com/yeli7068/VIP2 |
| virID | 2019 | reads (Illumina) | metaSPAdes | - | - | - | bwa mem | DIAMOND, MegaBLAST | - | https://github.com/jnoms/virID |
| phageannotator | NOT RELEASED YET | contigs + reads (Illumina) for abundance estimation only | - | geNomad, MASH | CheckV | VRhyme | bowtie2, CoverM | vConTACT3, geNomad | Prodigal-gv | https://github.com/nf-core/phageannotator/tree/dev |

**Supplementary Table 2. Detailed CPU time usage per job step.** Number of jobs run (either 1 common or 1 for each sample processed), the average and standard deviation of the CPU time used (in seconds) for the job, as well as the total CPU time used for each main step in the workflow for each dataset.

| | | Moreno-Gallego | | | | | Li | | | | | Buck | | | |
|---|---|---|---|---|---|---|---|---|---|---|---|---|---|---|---|
| **Part** | **Step** | **Jobs** | **Average** | **St.dev.** | **Sum** | | **Jobs** | **Average** | **St.dev.** | **Sum** | | **Jobs** | **Average** | **St.dev.** | **Sum** |
| **QC** | fastp_pe | 84 | 34 | 10 | 2 892 | | 196 | 312 | 49 | 61 234 | | 350 | 438 | 256 | 153 300 |
| **QC** | fastqc | 1 | 1 874 | 0 | 1 874 | | 1 | 41 740 | 0 | 41 740 | | 1 | 118 377 | 0 | 118 377 |
| **QC** | multiqc | 1 | 6 | 0 | 6 | | 1 | 20 | 0 | 20 | | 1 | 17 | 0 | 17 |
| **Asm** | metaSpades | 84 | 76 726 | 64 253 | 6 444 988 | | 196 | 69 049 | 12 280 | 13 533 526 | | 350 | 218 928 | 117 555 | 76 624 895 |
| **Asm** | metaQUAST | 1 | 25 445 | 0 | 25 445 | | 1 | 57 894 | 0 | 57 894 | | 1 | 107 282 | 0 | 107 282 |
| **ID** | virsorter | 84 | 1 953 | 1 028 | 164 081 | | 196 | 82 757 | 36 355 | 16 220 335 | | 350 | 100 221 | 78 680 | 35 077 455 |
| **ID** | checkv/virsorter | 84 | 75 | 22 | 6 287 | | 196 | 261 | 202 | 51 191 | | 350 | 1 614 | 2 007 | 564 806 |
| **ID** | virsorter_for_dram | 1 | 16 596 | 0 | 16 596 | | 1 | 93 585 | 0 | 93 585 | | 1 | 330 338 | 0 | 330 338 |
| **Map** | build_index | 1 | 8 | 0 | 8 | | 1 | 45 | 0 | 45 | | 1 | 223 | 0 | 223 |
| **Map** | bowtie2_mapping | 84 | 441 | 338 | 37 085 | | 196 | 1 660 | 319 | 325 295 | | 350 | 1 878 | 1 184 | 657 374 |
| **Map** | mapping/flagstat | 84 | 2 | 2 | 166 | | 196 | 35 | 5 | 6 791 | | 350 | 23 | 24 | 7 875 |
| **Map** | mapping/pileup | 84 | 13 | 4 | 1 110 | | 196 | 34 | 5 | 6 705 | | 350 | 36 | 11 | 12 751 |
| **Map** | instrain | 84 | 1 340 | 766 | 112 559 | | 196 | 2 357 | 772 | 461 970 | | 336 | 3 064 | 3 714 | 1 029 501 |
| **Map** | instrain/compare | 1 | 10 877 | 0 | 10 877 | | 1 | 449 436 | 0 | 449 436 | | 1 | 15 | 0 | 15 |
| **Tax** | prodigal | 1 | 47 | 0 | 47 | | 1 | 747 | 0 | 747 | | 1 | 1 927 | 0 | 1 927 |
| **Tax** | vcontact2_gene2genome | 1 | 1 | 0 | 1 | | 1 | 1 | 0 | 1 | | 1 | 4 | 0 | 4 |
| **Tax** | inphared_setup | 1 | 2 | 0 | 2 | | 1 | 1 | 0 | 1 | | 1 | 2 | 0 | 2 |
| **Tax** | vcontact2 | 1 | 72 111 | 0 | 72 111 | | 1 | 70 976 | 0 | 70 976 | | 1 | 86 261 | 0 | 86 261 |
| **Tax** | graphanalyzer | 1 | 179 | 0 | 179 | | 1 | 328 | 0 | 328 | | 1 | 2 522 | 0 | 2 522 |
| **Fun** | DRAMv | 1 | 9 963 | 0 | 9 963 | | 1 | 59 118 | 0 | 59 118 | | 1 | 217 160 | 0 | 217 160 |
| **Fun** | DRAMv_distill | 1 | 2 | 0 | 2 | | 1 | 1 | 0 | 1 | | 1 | 2 | 0 | 2 |

**Supplementary Table 3. Detailed wall time usage per job step.** Number of jobs run (either 1 common or 1 for each sample processed), the average and standard deviation of the wall time used (in seconds) for the job, as well as the total wall time used for each main step in the workflow for each dataset.

| | | Moreno-Gallego | | | | Li | | | | Buck | | | |
|---|---|---|---|---|---|---|---|---|---|---|---|---|---|
| **Part** | **Step** | **Jobs** | **Average** | **St.dev.** | **Sum** | **Jobs** | **Average** | **St.dev.** | **Sum** | **Jobs** | **Average** | **St.dev.** | **Sum** |
| **QC** | fastp_pe | 84 | 12 | 5 | 982 | 196 | 99 | 16 | 19 473 | 350 | 129 | 69 | 45 147 |
| **QC** | fastqc | 1 | 314 | 0 | 314 | 1 | 7 298 | 0 | 7 298 | 1 | 14 986 | 0 | 14 986 |
| **QC** | multiqc | 1 | 20 | 0 | 20 | 1 | 58 | 0 | 58 | 1 | 61 | 0 | 61 |
| **Asm** | metaSpades | 84 | 10 908 | 9 159 | 916 296 | 196 | 13 843 | 2 690 | 2 713 173 | 350 | 34 527 | 19 169 | 12 084 457 |
| **Asm** | metaQUAST | 1 | 5 175 | 0 | 5 175 | 1 | 15 425 | 0 | 15 425 | 1 | 27 706 | 0 | 27 706 |
| **ID** | virsorter | 84 | 833 | 232 | 70 002 | 196 | 18 152 | 6 410 | 3 557 872 | 350 | 15 317 | 11 086 | 5 360 828 |
| **ID** | checkv/virsorter | 84 | 44 | 16 | 3 713 | 196 | 196 | 84 | 38 422 | 350 | 391 | 365 | 136 973 |
| **ID** | virsorter_for_dram | 1 | 4 124 | 0 | 4 124 | 1 | 20 320 | 0 | 20 320 | 1 | 44 256 | 0 | 44 256 |
| **Map** | build_index | 1 | 8 | 0 | 8 | 1 | 63 | 0 | 63 | 1 | 251 | 0 | 251 |
| **Map** | bowtie2_mapping | 84 | 69 | 41 | 5 809 | 196 | 362 | 49 | 70 983 | 350 | 328 | 182 | 114 881 |
| **Map** | mapping/flagstat | 84 | 1 | 1 | 74 | 196 | 11 | 3 | 2 091 | 350 | 7 | 3 | 2 618 |
| **Map** | mapping/pileup | 84 | 4 | 1 | 333 | 196 | 24 | 4 | 4 747 | 350 | 22 | 8 | 7 595 |
| **Map** | instrain | 84 | 429 | 197 | 36 076 | 196 | 768 | 187 | 150 536 | 336 | 1 299 | 1 075 | 436 608 |
| **Map** | instrain/compare | 1 | 2 010 | 0 | 2 010 | 1 | 101 844 | 0 | 101 844 | 1 | 19 | 0 | 19 |
| **Tax** | prodigal | 1 | 61 | 0 | 61 | 1 | 759 | 0 | 759 | 1 | 1 957 | 0 | 1 957 |
| **Tax** | vcontact2_gene2genome | 1 | 5 | 0 | 5 | 1 | 6 | 0 | 6 | 1 | 7 | 0 | 7 |
| **Tax** | inphared_setup | 1 | 5 | 0 | 5 | 1 | 6 | 0 | 6 | 1 | 6 | 0 | 6 |
| **Tax** | vcontact2 | 1 | 25 783 | 0 | 25 783 | 1 | 29 658 | 0 | 29 658 | 1 | 36 767 | 0 | 36 767 |
| **Tax** | graphanalyzer | 1 | 144 | 0 | 144 | 1 | 323 | 0 | 323 | 1 | 1 055 | 0 | 1 055 |
| **Fun** | DRAMv | 1 | 3 502 | 0 | 3 502 | 1 | 19 893 | 0 | 19 893 | 1 | 72 979 | 0 | 72 979 |
| **Fun** | DRAMv_distill | 1 | 17 | 0 | 17 | 1 | 26 | 0 | 26 | 1 | 28 | 0 | 28 |
